# The client-binding domain of the cochaperone Sgt2 has a helical-hand structure that binds a short hydrophobic helix

**DOI:** 10.1101/517573

**Authors:** Ku-Feng Lin, Michelle Y. Fry, Shyam M. Saladi, William M. Clemons

## Abstract

The targeting and insertion of tail-anchored (TA) integral membrane proteins (IMP) into the correct membrane is critical for cellular homeostasis. The fungal protein Sgt2, and its human homolog SGTA, binds hydrophobic clients and is the entry point for targeting of ER-bound TA IMPs. Here we reveal molecular details that underlie the mechanism of Sgt2 binding to TA IMP clients. We establish that the Sgt2 C-terminal region is flexible but conserved and sufficient for client binding. A molecular model for this domain reveals a helical hand forming a hydrophobic groove, consistent with a higher affinity for TA IMP clients with hydrophobic faces and a minimal length of 11 residues. This work places Sgt2 into a broader family of TPR-containing co-chaperone proteins.

## Introduction

An inherently complicated problem of cellular homeostasis is the biogenesis of hydrophobic IMPs which are synthesized in the cytoplasm and must be targeted and inserted into a lipid bilayer. Accounting for ~25% of transcribed genes [1], IMPs are primarily targeted by cellular signal binding factors that recognize a diverse set of hydrophobic α-helical signals as they emerge from the ribosome [2–4]. One important class of IMPs are tail-anchored (TA) proteins whose hydrophobic signals are their single helical transmembrane domain (TMD) located near the C-terminus and are targeted post-translationally to either the ER or mitochondria [5–9]. In the case of the canonical pathway for ER-destined TA IMPs, each is first recognized by homologs of mammalian SGTA (small glutamine tetratricopeptide repeat protein) [4,6,10,11]. Common to all signal binding factors is the need to recognize, bind, and then hand off a hydrophobic helix. How such factors can maintain specificity to a diverse set of hydrophobic clients that must subsequently be released remains an important question.

Homologs of *Saccharomyces cerevisiae* Sgt2 (ySgt2) and *Homo sapiens* SGTA (referred to here as hSgt2 for simplicity), collectively Sgt2, are involved in a variety of cellular processes regarding the homeostasis of membrane proteins including the targeting of TA IMPs [9,12–14], retrograde transport of membrane proteins for ubiquitination and subsequent proteasomal degradation [15], and regulation of mislocalized membrane proteins (MLPs) [16,17]. Among these, the role of Sgt2 in the primary pathways responsible for targeting TA clients to the endoplasmic reticulum (ER) is best characterized, *i.e.* the fungal <u>G</u>uided <u>E</u>ntry of <u>T</u>ail-anchored proteins (GET) or the mammalian <u>T</u>ransmembrane <u>R</u>ecognition <u>C</u>omplex (TRC) pathway. In the GET pathway, Sgt2 functions by binding a cytosolic TA client then transferring the TA client to the ATPase chaperone Get3 (human homolog is also Get3) with the aid of the heteromeric Get4/Get5 complex (human Get4/Get5/Bag6 complex) [13,18–20]. In this process, TA client binding to Sgt2, after hand-off from Hsp70, is proposed as the first committed step to ensure that ER TA clients are delivered to the ER membrane while mitochondrial TA clients are excluded [3,13,21]. Subsequent transfer of the TA client from Sgt2 to the ATP bound Get3 induces conformational changes in Get3 that trigger ATP hydrolysis, releasing Get3 from Get4 and favoring binding of the Get3-TA client complex to the Get1/2 (human Get1/Get2) receptor at the ER leading to release of the TA client into the membrane [22–26]. Deletions of yeast GET genes (*i.e. get1*Δ, *get2*Δ, or *get3*Δ) cause cytosolic aggregation of TA clients dependent on Sgt2 [26,27].

In addition to targeting TA IMPs, there is evidence hSgt2 promotes degradation of IMPs through the proteasome by cooperating with the Bag6 complex, a heterotrimer containing Bag6, hGet4, and hGet5, which acts as a central hub for a diverse physiological network related to protein targeting and quality control [19,28–30]. The Bag6 complex can associate with ER membrane-embedded ubiquitin regulatory protein UbxD8, transmembrane protein gp78, proteasomal component Rpn10c, and an E3 ubiquitin protein ligase RNF126 thereby connecting hSgt2 to ER associated degradation (ERAD) and proteasomal activity. Depletion of hSgt2 significantly inhibits turnover of ERAD IMP clients and elicits the unfolded protein response[16]. Furthermore, the cellular level of MLPs in the cytoplasm could be maintained by co-expression with hSgt2, which possibly antagonize ubiquitination of MLPs to prevent proteasomal degradation [15,17]. These studies demonstrate an active role of hSgt2 in triaging IMPs in the cytoplasm and the breadth of hSgt2 clients including TA IMPs, ERAD, and MLPs all harboring one or more TMD. Roles for hSgt2 in disease include polyomavirus infection [31], neurodegenerative disease [27,32], hormone-regulated carcinogenesis [33,34], and myogenesis [35], although the underlying molecular mechanisms are still unclear.

The architecture of Sgt2 includes three structurally independent domains that define the three different interactions of Sgt2 (Fig. 1*A*) [12,36–39]. The N-terminal domain forms a homo-dimer composed of a four-helix bundle with 2-fold symmetry that primarily binds to the ubiquitin-like domain (UBL) of Get5/Ubl4A for TA IMP targeting [36,40] or interacts with the UBL on the N-terminal region of BAG6 [41] where it is thought to initiate downstream degradation processes [15,28,29]. The central region comprises a co-chaperone domain with three repeated TPR motifs arranged in a right handed-superhelix forming a ‘carboxylate clamp’ for binding the C-terminus of heat-shock proteins (HSP) [12,42]. The highly conserved TPR domain was demonstrated to be critical in modulating propagation of yeast prions by recruiting HSP70 [27] and may associate with the proteasomal factor Rpn13 to regulate MLPs [43]. More recently, it was demonstrated that mutations to residues in the TPR domain which prevent Hsp70 binding impair the loading of TA clients onto ySgt2 [21], consistent with a direct role of Hsp70 in TA IMP targeting via the TPR domain. The C-terminal methionine-rich domain of Sgt2 is responsible for binding to hydrophobic clients such as TA IMPs [11,37,44]. Other hydrophobic segments have been demonstrated to interact with this domain such as the membrane protein Vpu (viral protein U) from human immunodeficiency virus type-1 (HIV1), the TMD of tetherin [44], the signal peptide of myostatin [35], and the N-domain of the yeast prion forming protein Sup35 [27]. All of these studies suggest that the C-terminus of Sgt2 binds broadly to hydrophobic stretches, yet structural and mechanistic information for client recognition is lacking.

**Fig. 1.**
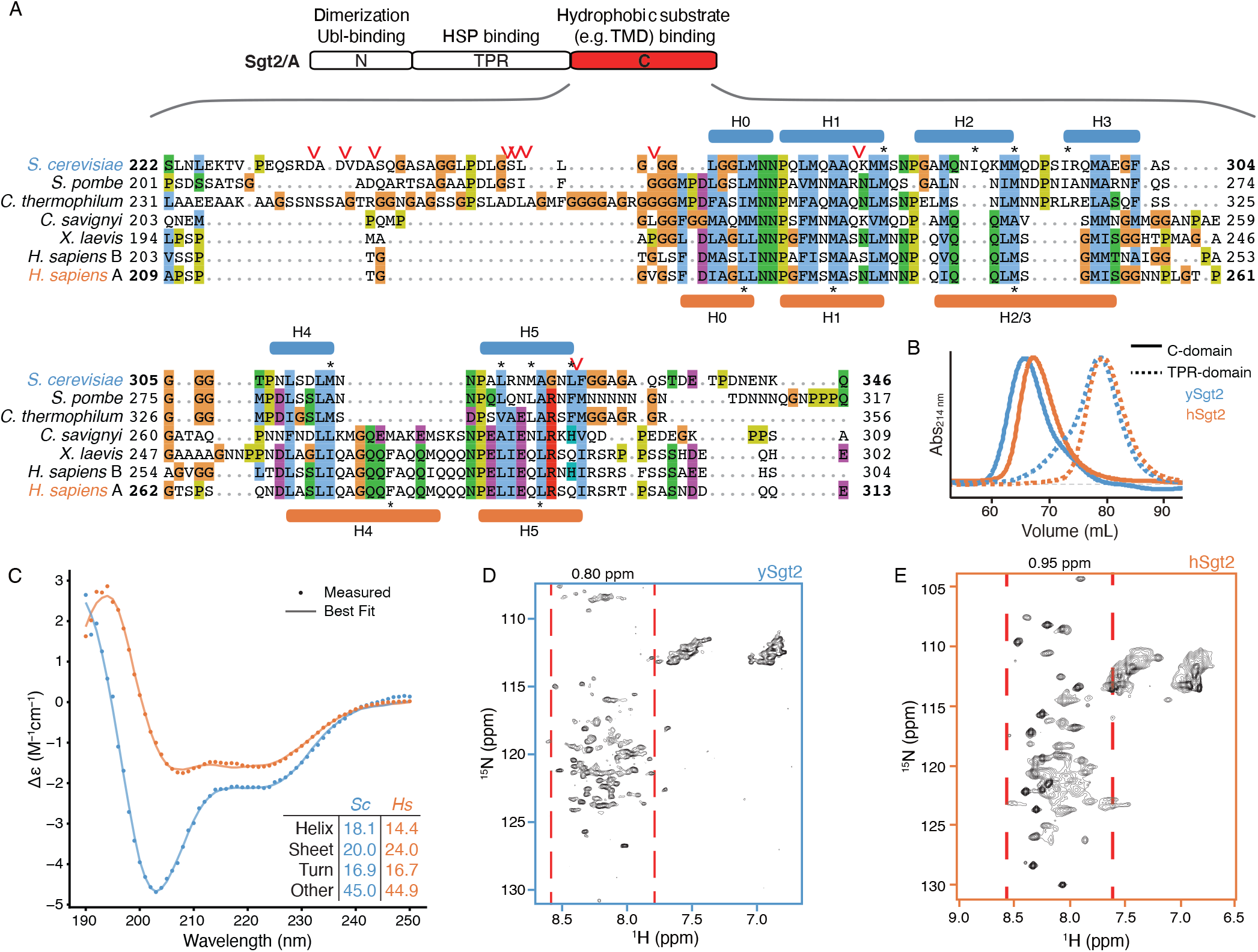
Structural characteristics of free Sgt2 C-domain. (*A*) *Top*, Schematic of the domain organization of Sgt2. *Below,* representative sequences from a large-scale multiple sequence alignment of the C domain: fungal Sgt2 from *S. cerevisiae*, *S. pombe*, and *C. thermophilum* and metazoan Sgt2 from *C. savignyi*, *X. laevis*, and *H. sapiens*. Protease susceptible sites on ySgt2-C identified by mass spectrometry are indicated by red arrowheads. Predicted helices of ySgt2 (blue) and hSgt2 (orange) by Jpred [83] and/or structure prediction are shown. Blue/orange color scheme for ySgt2/hSgt2 is used throughout the text. Residues noted in the text are highlighted by an asterisk. (*B*) Overlay of size-exclusion chromatography traces of ySgt2-C (blue line), hSgt2-C (orange line), ySgt2-TPR (blue dash), and hSgt2-TPR (orange dash). Traces are measured at 214 nm, baseline-corrected and normalized to the same peak height. (*C*) Far UV CD spectrum of 10 μM of purified ySgt2-C (blue) and hSgt2-C (orange) at RT with secondary structure decomposition from BestSel [68]. (*D*) _1_H-_15_N HSQC spectrum of ySgt2-C at 25°C. The displayed chemical shift window encompasses all N-H resonances from both backbone and side chains. The range of backbone amide protons, excluding possible side-chain NH2 of Asn/Gln, is indicated by pairs of red dashed lines. (*E*) As in *D* for hSgt2-C at 25°C.

In this study, we provide the first structural characterization of the C-domains from Sgt2 (Sgt2-C) and show that, in the absence of substrate, it is relatively unstructured. We demonstrate that a conserved region of the C-domain, defined here as C_cons_, is sufficient for client binding. Analysis of the C_cons_ sequence identifies six amphipathic helices whose hydrophobic residues are required for client binding. Based on this, we computationally generate an *ab initio* structural model that is validated by point mutants and disulfide crosslinking. Artificial TA clients are then used to define the properties within clients critical for binding to Sgt2-C. The results show that Sgt2-C falls into a larger STI1 family of TPR-containing co-chaperones and allow us to propose a mechanism for TA client binding.

## Results

### The flexible Sgt2-C domain

Based on sequence alignment (Fig. 1*A*), the Sgt2-C contains a conserved core of six predicted helices flanked by unstructured loops that vary in length and sequence. Previous experimental work suggested that this region is particularly flexible, as this domain in the *Aspergillus fumigatus* is sensitive to proteolysis [12]. Similarly, for ySgt2-TPR-C, the sites sensitive to limited proteolysis primarily occur within the loops flanking the conserved helices (Fig. 1*A*, *red arrows* and S1*B*). This flexible nature of the C-domain likely contributes to its anomalous passage through a gel-filtration column where Sgt2-C elutes much earlier than the similarly-sized, but well-folded Sgt2 TPR-domain (Fig. 1*B*). The larger hydrodynamic radius matches previous small-angle X-ray scattering measurement of the ySgt2 TPR-C domain that indicated a partial unfolded characteristic in a Kratky plot analysis. The circular dichroism (CD) spectra for both homologs suggests that the C-domain largely assumes a random-coil conformation, with 40-45% not assignable to a defined secondary structure category (Fig. 1*C*) [45]. The well-resolved, sharp, but narrowly dispersed chemical shifts of the backbone amide protons in 1H-15N HSQC spectra of Sgt2-C (Fig. 1*D,E*), indicate a significant degree of backbone mobility, similar to natively unfolded proteins [46] and consistent with results seen by others [47], further highlighting the lack of stable tertiary structure. [12]. Taken all together, Sgt2-C appears to be a flexible domain.

### The conserved region of the C-domain is sufficient for substrate binding

We then asked if the flexible Sgt2-C is the site of client binding in the co-chaperone and if so, where within this domain is the binding region. During purification Sgt2-C is susceptible to proteolytic activity being cut at several specific sites (Fig. 1*A*). Proteolysis occurred primarily at Leu_327_ and in the poorly conserved N-terminal region (between Asp_235_-Gly_258_). Given the intervening region, between Gly_258_ and Leu_327_ on ySgt2, is conserved (Fig 1*A*), it and the corresponding region on hSgt2, may mediate TA client binding (Fig. 2*A*, *grey*). To test this, we established a set of his-tagged Sgt2 constructs of various lengths (Fig. 2*C*). These Sgt2-C mutants were co-expressed with an MBP-tagged TA client, Sbh1, and binding was detected by the presence of captured TA clients in nickel elution fractions (Fig. 2*B*). As previously seen [13], we confirm that Sgt2-TPR-C alone is sufficient for capturing a TA client (Fig. 2*C*). As one might expect, the C-domain was also sufficient for binding the TA client. A predicted six α-helical methionine-rich region of Sgt2-C (Fig. 1*A*), hereafter referred to as Sgt2-C_cons_, is sufficient for binding to Sbh1. For ySgt2, a minimal region H1-H5 (ΔH0) poorly captures Sbh1, while for hSgt2 the equivalent minimal region is sufficient for capturing the client at a similar level as the longer C_cons_ domain (Fig. 2*C*). The predicted helices in Sgt2-C_cons_ are amphipathic and their hydrophobic patches could be used for client binding (Fig. 2*D*).

**Fig. 2.**
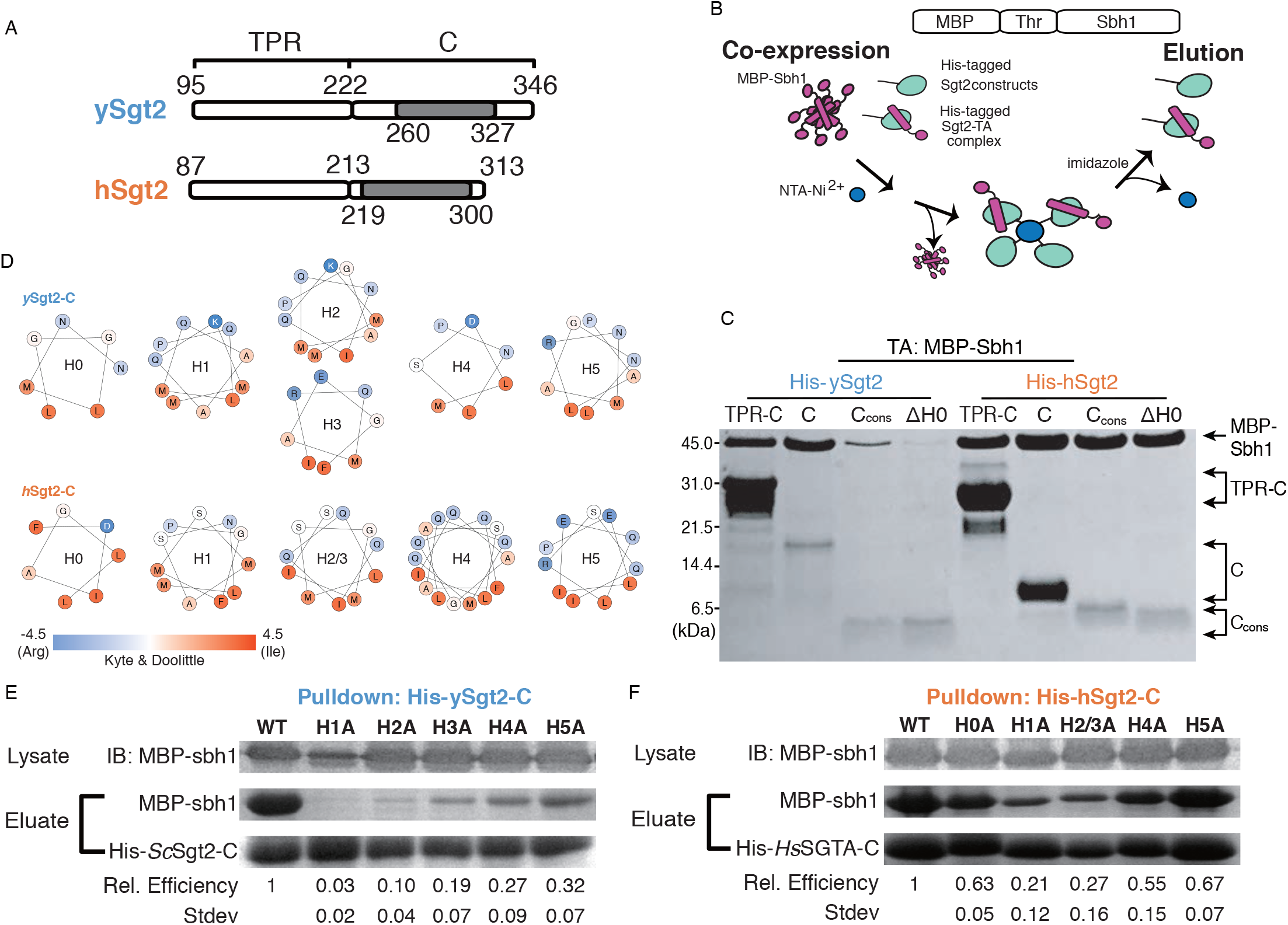
The minimal binding region of Sgt2 for TA client binding. (*A*) Diagram of the protein truncations tested for TA client binding that include the TPR-C domain, C-domain (C), C_cons_, and C_cons_ ΔH0 (ΔH0) from ySgt2 and hSgt2. The residues corresponding to each domain are indicated, and grey blocks highlight the C_cons_ region. *(B)* Schematic of capture experiments of MBP-tagged Sbh1 (MBP-Sbh1) by Sgt2 variants. After co-expression, cell pellets are lysed and NTA-Ni_2+_ is used to capture his-tagged Sgt2-TPR-C. (*C*) Tris-Tricine-SDS-PAGE gel [84] of co-expressed and purified MBP-Sbh1 and his-tagged Sgt2 truncations visualized with Coomassie Blue staining. (*D*) Helical wheel diagrams of predicted helices (see Fig. 1*A*) in the C_cons_ domain of ySgt2 and hSgt2. Residues are colored by the Kyte and Doolittle hydrophobicity scale [85]. (*E*) All of the hydrophobic residues (L, I, F, and M) in a predicted helix (H0, H1, etc.) are replaced with alanines and tested for the ability to capture MBP-Sbh1. Protein levels were quantified by Coomassie staining. Relative binding efficiency of MBP-Sbh1 by ySgt2 C-domain (ySgt2-C) variants was calculated relative to total amount of ySgt2-C captured (MBP-Sbh1/Sgt2-C) then normalized to the wild-type ySgt2-C. Experiments were performed 3-4 times and the standard deviations are presented. Total expression levels of the MBP-Sbh1 were similar across experiments as visualized by immunoblotting (IB) of the cell lysate. (*F)* As in *E* but for hSgt2.

To test this, each of the six helices in Sgt2-C_cons_ was mutated to replace the larger hydrophobic residues with alanines, dramatically reducing the overall hydrophobicity. For all of the helices, alanine replacement of the hydrophobic residues significantly reduces binding of Sbh1 to Sgt2-C (*Fig. 2E & F*). While these mutants expressed at similar levels to the wild-type sequence, one cannot rule out that some of these changes may affect the tertiary structure of this domain. In general, these results imply that these amphipathic helices are necessary for client binding since removal of the hydrophobic faces disrupts binding. The overall effect on binding by each helix is different, with mutations in helices 1-3 having the most dramatic reduction in binding suggesting that these are more crucial for Sgt2-TA client (Sgt2-TA) complex formation. It is also worth noting, as this is a general trend, that hSgt2 is more resistant to mutations that affect binding (Fig. 2*F*) than ySgt2, which likely reflect different thresholds for binding.

### Molecular modeling of Sgt2-C domain

Despite the need for a molecular model, the C-domain has resisted structural studies, likely due to the demonstrated inherent flexibility. Based on the six conserved α-helical amphipathic segments (Fig. 1*A*) that contain hydrophobic residues critical for TA client binding (Fig. 2*D-E*), we expect some folded structure to exist. Therefore, we performed *ab initio* molecular modeling of Sgt2-C using a variety of prediction methods resulting in a diversity of putative structures [48–52]. As expected, all models showed buried hydrophobic residues as this is a major criterion for *in silico* protein folding. Residues outside the ySgt2-C_cons_ region adopted varied conformations consistent with their expected higher flexibility. Pruning these N- and C-terminal regions to focus on the ySgt2-C_cons_ region (Fig. S2*A*) revealed a potential binding interface for a hydrophobic substrate, examples are seen in Quark models (1, 4, & 6 shown), Robetta 1 & 2, and I-TASSER 2 & 3, whereas others models had no clearly distinguishable groove. Given the intrinsic flexibility of the Sgt2-C domain, it is possible that models without a groove are found in the non-TMD bound structural ensemble.

For a working model of TMD-bound ySgt2-C, we chose the highest scored Quark structures where a general consistent architecture was seen (Fig. 3*A*) [48]. The overall model contained a potential TA client binding site, a hydrophobic groove formed by the amphipathic helices. The groove is approximately 15 Å long, 12 Å wide, and 10 Å deep, which is sufficient to accommodate three helical turns of an α-helix, ~11 amino acids (Fig. 3*B*).

**Fig 3.**
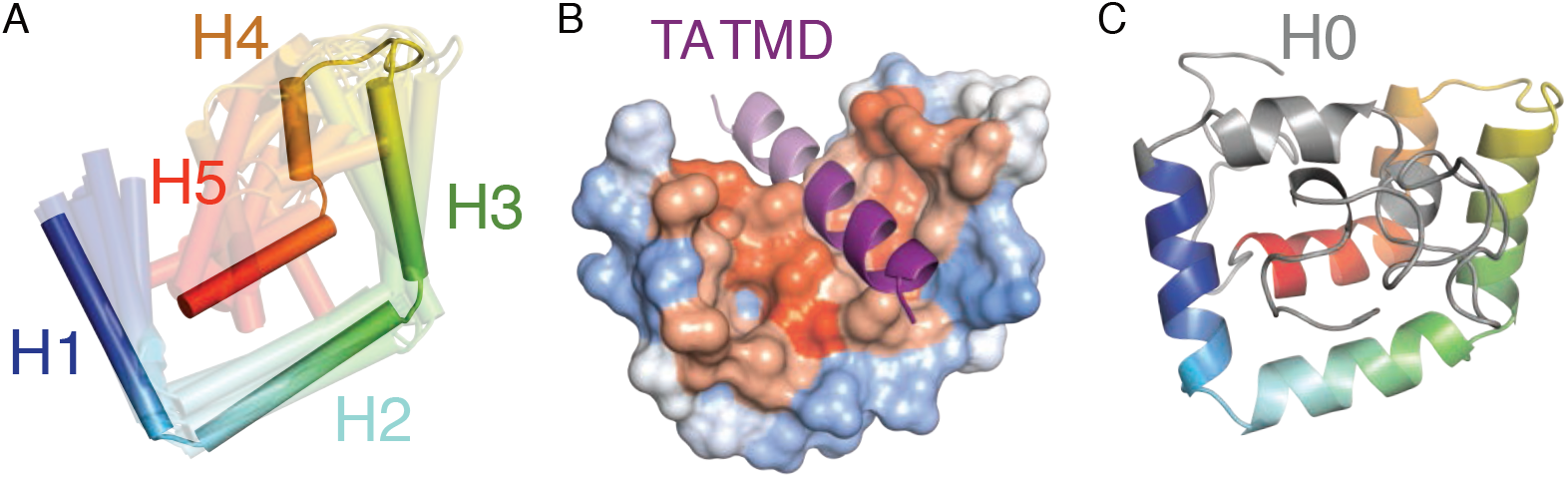
A structural model for Sgt2-C_cons_. *(A)* The top 10 models of the ySgt2-C_cons_ generated by the template-free algorithm Quark [48] are overlaid with the highest scoring model in solid. Models are color-ramped from N-(blue) to C-terminus (red). *(B)* A model of ySgt2-C_cons_ (surface colored by Kyte-Doolittle hydrophobicity) bound to a TMD (purple helix) generated by rigid-body docking through Zdock [80]. The darker purple corresponds to an 11 residue stretch. *(C)* The entire ySgt2-C from the highest scoring model from Quark (C_cons_ in rainbow with the rest in grey) highlighting H0 and the rest of the flexible termini that vary considerably across models.

To validate the model, we interrogated the accuracy of the predicted structural arrangement by determining distance constraints from crosslinking experiments. We selected four pairs of residues in close spatial proximity and one pair far apart based on the Quark models (Fig. 4*A*). Calculating a C_β_-C_β_ distance between residue pairs for each model (Fig. 4*E*), the Quark models 2 and 3 were the most consistent with an expected distance of 9Å or less for the close pairs. In all alternative models, the overall distances are much larger and should not be expected to form disulfide bonds *in vitro* if they represent a TMD-bound state. For Robetta, a number of the models have pairs of residues within 9Å and Robetta’s per-residue error estimate suggests relatively high confidence in the C_cons_ region (Fig. S2*B*).

**Fig 4.**
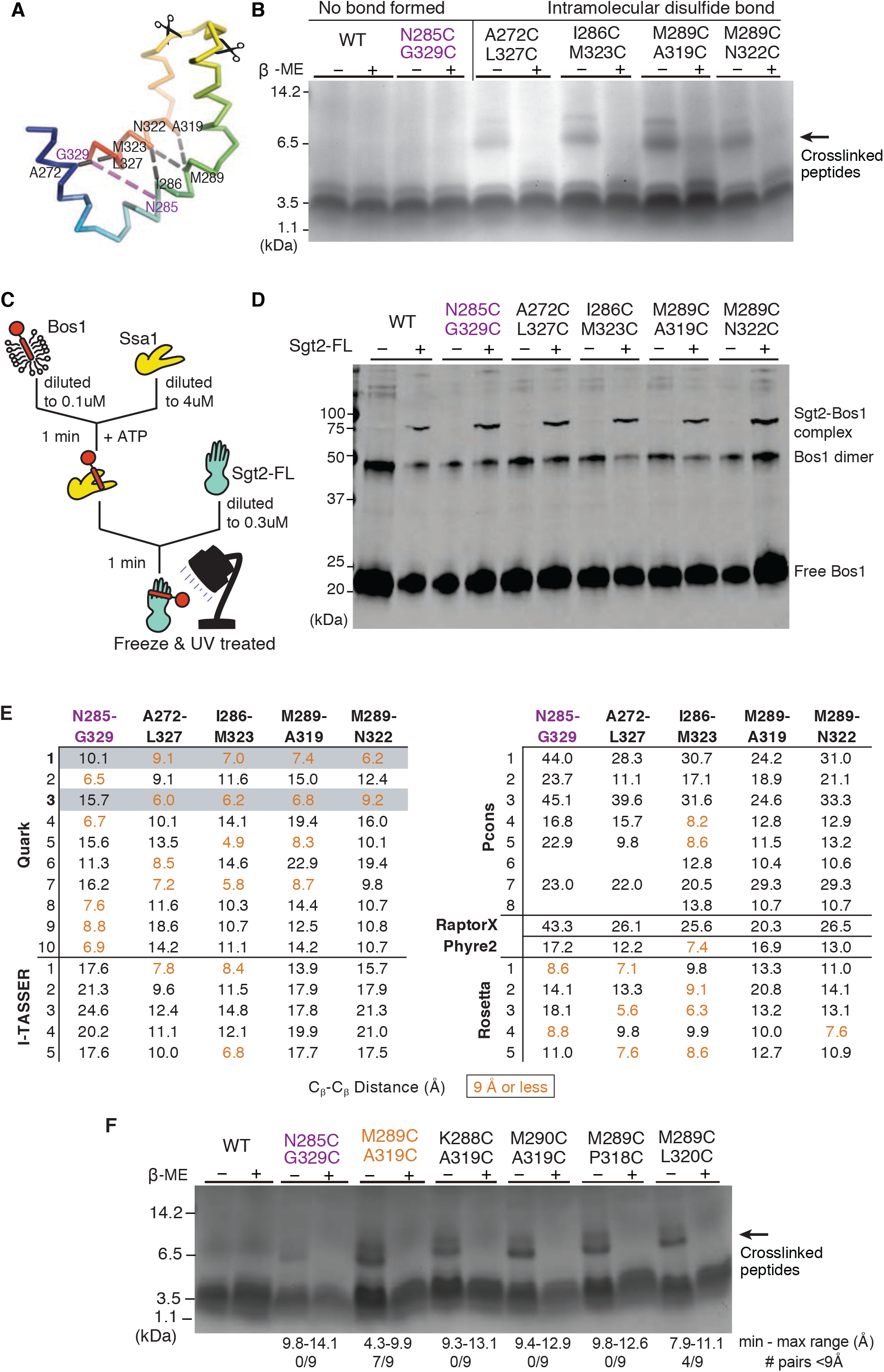
Validating the structural model with disulfide bond formation. Variants of His-ySgt2-TPR-C (WT or cysteine double mutants) were co-expressed with the artificial TA client, cMyc-BRIL-11[L8]. After lysis, ySgt2-TPR-C proteins were purified, oxidized, then digested by Glu-C protease and analyzed by gel either in non-reducing or reducing buffer. (*A*) Cα ribbon of ySgt2-C_cons_ color-ramped with various pairs of cysteines highlighted. Scissors indicate protease cleavage sites resulting in fragments less than 3 kDa in size. (*B*) Tris-Glycine-SDS-PAGE gel visualized by imidazole-SDS-zinc stain [86]. For the WT (cys-free) no significant difference was found between samples in non-reducing vs. reducing conditions. All close residue pairs (A272/L327, I286/M323, M289/A319, and M289/N322) show peptide fragments (higher MW) sensitive to the reducing agent and indicate disulfide bond formation (indicated by arrow). A cysteine pair (N285/G329) predicted to be far apart by the model does not result in the higher MW species. (*C*) A schematic of the transfer of Bos1_BPA_ from Ssa1 to full-length ySgt2 to demonstrate the double cysteine mutants are still functional. (*D*) A western blot visualizing cross-linked ySgt2-Bos1_BPA_ complexes. All samples tested, WT, N285C/G329C, A272C/L327C, I286C/M323C, M289C/A319C, and M289C/N322C, had a higher molecular weight appear after the addition of ySgt2 which corresponds to the size of the cross-linked complex. As described in Cho et al., a band for cross-linked Ssa1-Bos1_BPA_ complexes was not observed. (*E*) C_β_-C_β_ distances between the residues mutated to cysteines based on various models predicted by the Quark, I-TASSER, PCONS, and Robetta. Cysteine pairs that are 9Å or less apart are colored in orange and are expected to be close enough to form disulfide bonds. Where all five pair distances are consistent with the experiment (4 near and 1 far), the row is shaded in grey. (*F*) Tris-Glycine SDS-PAGE gel probing the flexibility of ySgt2-C_cons_. All new pairs (K288/A319, M290/A319, M289/P318, M289/L320) show peptide fragments sensitive to the reducing agent (indicated by arrow). The range of distances of the eight closest possible rotamer pairs is annotated below. The cysteine pair (N285/G329) shown to be far apart by the model does have a faint higher molecular weight band.

As a control, we first confirmed that the cysteine-mutant pairs do not affect the function of ySgt2. We utilized an *in vitro* transfer assay where a yeast Hsp70 homolog Ssa1 loaded with a TA client delivers the client to ySgt2 [21,49,50] (Fig. 4*C*). Purified Ssa1 is mixed with detergent solubilized strep-tagged Bos1-TMD (a model ER TA client) that contained a p-benzoyl-l-phenylalanine (BPA) labeled residue, Bos1_BPA_, are diluted to below the critical micelle concentration resulting in soluble complexes of Bos1_BPA_/Ssa1. Full-length ySgt2 variants were each tested for the ability to capture Bos1_BPA_ from Ssa1. After the transfer reaction, each was UV-treated to generate Bos1 crosslinks. Successful capture of the TA clients by ySgt2 was detected using an anti-strep Western blot and the appearance of a Bos1_BPA_/ySgt2 crosslink band (Fig. 4*D*). All of the cysteine variants of ySgt2 successfully captured Bos1BPA from Ssa1 similar to wild-type suggesting that the cysteine mutations did not affect the structure or function of ySgt2.

For the distance experiment, each of the cysteine-mutant pairs was made in ySgt2-TPR-C which lacks the dimerization domain. Each variant was coexpressed with an artificial TA client, a cMyc-tagged BRIL (small, 4-helix bundle protein [51]) with a C-terminal TMD consisting of eight leucines and three alanines, denoted 11[L8], and purified via nickel-affinity chromatography in reducing buffer (Fig. S3*A*). All of the ySgt2 mutants bound the TA client and behaved similar to the wild-type (cysteine-free) further suggesting the mutants did not perturb the native structure (Fig. S3*B*). For disulfide crosslink formation, each eluate was oxidized and crosslinks were identified by the visualization of a reducing-agent sensitive ~7.7kDa fragment in gel electrophoresis (Fig. 3*B*). For both the wild-type construct and in N285C/G329C, where the pairs are predicted from the Quark models to be too distant for disulfide bond formation, no higher molecular weight band was observed. For the remaining pairs that are predicted to be close enough for bond formation, the 7.7kDa fragment was observed in each case and is labile in reducing conditions. Again, these results support the C_cons_ model derived from Quark.

With the four crosslinked pairs as distance constraints, new models were generated using Robetta with a restraint on the corresponding pairs of C_β_ atoms less than 9Å (Fig. S4*A*). The Robetta models from these runs are similar to the top scoring models from Quark (Fig. 3). Satisfyingly, the pair of residues that do not form disulfide crosslinks are generally consistent (Fig. S4*B*).

The improvement of the ySgt2 models predicted by Robetta with restraints included encouraged us to generate models for hSgt2-C with constraints. For this, pairs were defined based on sequence alignments of Sgt2 (Fig. 1*A*) and used as restraints. The resulting predictions had architectures consistent with the equivalent regions predicted for ySgt2-C_cons_, for example Robetta 4 (Fig. S4*C*, top). Although in general the predicted hSgt2 model is similar to that for ySgt2, the region that corresponds to H2 occupies a position that precludes a clear hydrophobic groove. For ySgt2, the longer N-terminal loop occupies the groove preventing the exposure of hydrophobics to solvent (Fig. 3*C*, grey). For hSgt2, the shorter N-terminal loop may not be sufficient to similarly occupy the groove and allowing for the clear hydrophobic hand seen for the ySgt2-C. To correct for this, we replaced the sequence of the N-terminal loop of hSgt2-C with the ySgt2-C loop and ran structure prediction with the pairwise distance restraints. This resulted in a model where the loop occupies the groove and, when pruned away suggests the hydrophobic hand seen in yeast (Fig. S4*C*, middle boxed). Of note, we also generated models of hSgt2-C using the most recent Robetta method (transform-restrained) which produces new structures with a groove and similar helical-hand architecture across the board (Fig. S4*C*, bottom).

We sought to further test the robustness of our model considering the intrinsic flexibility of Sgt2-C by probing for disulfide bond formation with neighboring residues of one of our crosslinking pairs. While the C_β_-C_β_ distance puts these adjacent pairs at farther than 9Å, mutating residues to cystines and measuring S-S distances across all possible pairs of rotamers provides a wider interval on possible distances and, therefore, the likelihood a disulfide bond will form (Fig. 4*F*). Cysteine mutants were introduced to the residues adjacent to M289 and A319 in ySgt2-TPR-C for four additional pairs: K288C/A319C, M290C/A319C, M289C/P318C, and M289C/L320C. As described previously, these mutants were coexpressed with a TA substrate, in this case the artificial BRIL-11[L8] which has a MBP-tag instead of a cMyc-tag. Complexes were purified by amylose and then nickel affinity chromatography to ensure eluates contained only Sgt2-TPR-C bound to substrate. Eluates were incubated in oxidizing conditions, quenched with 50mM NEM, and digested with Glu-C protease. Again, a reductant sensitive band at 7.7kDa is observed for each of these adjacent pairs. While the geometry of each of these C-C pairs might suggest against disulfide bond formation, given the intrinsic flexibility of Sgt2-C, it is not surprising that each of these pairs are able to form disulfide bonds. As before, disulfide bond formation was detected for the M289C/A319C pair. In this new construct, we now see a small amount of disulfide bond formation in the distant N285C/G329C pair, likely an effect of switching to the MBP tag.

### Structural similarity of Sgt2-C domain to STI1 domains

Attempts to glean functional insight for Sgt2-C from BLAST searches did not reliably return other families or non-Sgt2 homologs making functional comparisons difficult. A more extensive profile-based search using hidden Markov models from the SMART database [52] identified a similarity to domains in the yeast co-chaperone Sti1 (HOP in mammals). First called DP1 and DP2, due to their prevalence of aspartates (D) and prolines (P), these domains have been shown to be required for client-binding by Sti1 [53,54] and are termed ‘STI1’-domains in bioinformatics databases [52]. In yeast Sti1 and its human homolog HOP (combined will be referred to here as Sti1), each of the two STI1 domains (DP1 and DP2) are preceded by Hsp70/90-binding TPR domains, similar to the domain architecture of Sgt2. Deletion of the second, C-terminal STI1-domain (DP2) from Sti1 *in vivo* is detrimental, impairing native activity of the glucocorticoid receptor [53]. *In vitro*, removal of the DP2 domain from Sti1 results in the loss of recruitment of the progesterone receptor to Hsp90 without interfering in Sti1-Hsp90 binding [55]. These results implicate DP2 in binding of Sti1 clients. In addition, others have noted that, broadly, STI1-domains may present a hydrophobic groove for binding the hydrophobic segments of a client [53,54]. Furthermore, the similar domain organizations (*i.e.* Sgt2 TPR-C, Sti1 TPR-STI1) and molecular roles could imply an evolutionary relationship between these co-chaperones. Indeed, a multiple sequence alignment of the Sgt2-C_cons_ with several yeast STI1 domains (Fig. 5*A*) reveals strong conservation of structural features. H1-H5 of the predicted helical regions in C_cons_ align directly with the structurally determined helices in the DP2 domain of Sti1; this includes complete conservation of helix breaking prolines and close alignment of hydrophobic residues in the amphipathic helices [53].

**Fig. 5.**
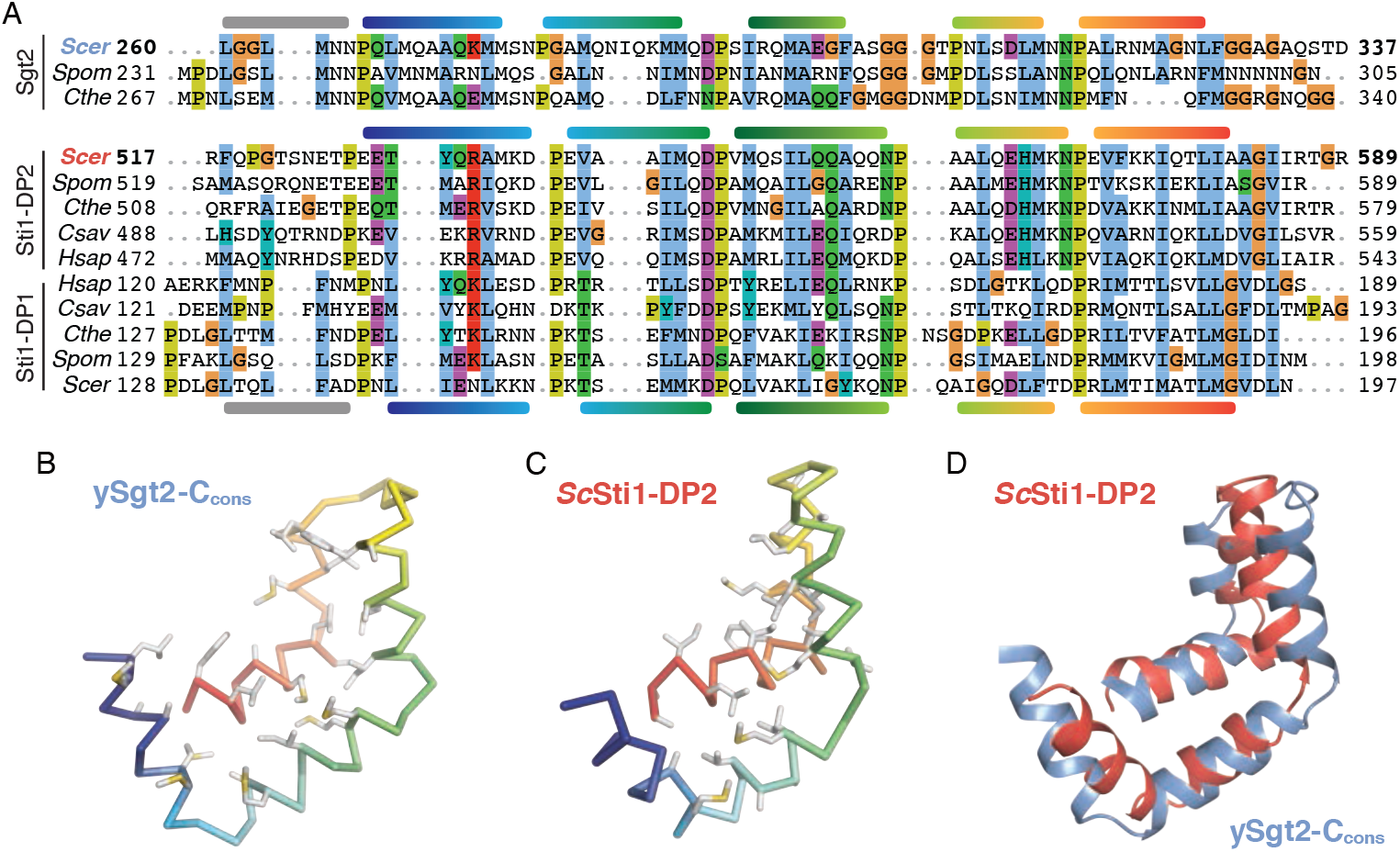
Comparison of Sti1 domains and the Sgt2-C_cons_ model. (*A*) Multiple sequence alignment of Sgt2-C with STI1 domains (DP1, DP2) from STI1/Hop homologs. Helices are shown based on the Sgt2-C_cons_ model and the *Sc*Sti1-DP1/2 structures. Species for representative sequences are from *S. cerevisiae (Scer)*, *S. pombe* (*Spom*), *C. thermophilum* (*Cthe*), *C. savignyi* (*Csav*), and *H. sapiens* (*Hsap*). (*B*) Cα ribbon of *Sc*Sgt2-C_cons_ color-ramped with large hydrophobic sidechains shown as grey sticks (sulfurs in yellow). (*C*) Similar to *B* for the solution NMR structure of Sti1-DP2_526-582_ (PDBID: 2LLW) [53]. (*D*) Superposition of the Sgt2-C_cons_ (blue) and Sti1-DP2_526-582_ (red) drawn as cartoons.

Based on the domain architecture and homology, a direct comparison between the STI1 domain and Sgt2-C_cons_ can be made. A structure of DP2 solved by solution NMR reveals that the five amphipathic helices assemble to form a flexible helical-hand with a hydrophobic groove [53]. The lengths of the α-helices in this structure concur with those inferred from the alignment in Fig. 4*A*. Our molecular model of Sgt2-C_cons_ is strikingly similar to this DP2 structure (Fig. 5*B,C*). An overlay of the DP2 structure and our molecular model demonstrates both Sgt2-C_cons_ and DP2 have similar lengths and arrangements of their amphipathic helices (Fig. 5*D*). Consistent with our observations of flexibility in Sgt2-C_cons_, Sti1-DP2 generates few long-range NOEs between its helices indicating that Sti1-DP2 also has a flexible architecture [53]. We consider this flexibility a feature of these helical-hands for reversible and specific binding of a variety of clients.

### Binding mode of TA clients to Sgt2

We examined the Sgt2-C_cons_ surface that putatively interacts with TA clients by constructing hydrophobic-to-charge residue mutations that are expected to disrupt capture of TA clients by Sgt2. Similar to the helix mutations in Fig. 2, the capture assay was employed to establish the relative effects of individual mutations. A baseline was established based on the amount of the TA client Sbh1 captured by wild-type Sgt2-TPR-C. In each experiment, Sbh1 was expressed at the same level; therefore, differences in binding should directly reflect the affinity of Sgt2 mutants for clients. In all cases, groove mutations from hydrophobic to aspartate led to a reduction in TA client binding (Fig. 6). The effects are most dramatic with ySgt2 where each mutant significantly reduced binding by 60% or more (Fig. 6*A*). While all hSgt2 individual mutants saw a significant loss in binding, the results were more subtle with the strongest a ~36% reduction (M233D, Fig. 6*B*). Double mutants were stronger with a significant decrease in binding relative to the individual mutants, more reflective of the individual mutants in ySgt2. As seen before (Fig. 2*E**&F*), we observe that mutations toward the N-terminus of Sgt2-C have a stronger effect on binding than those later in the sequence.

**Fig. 6.**
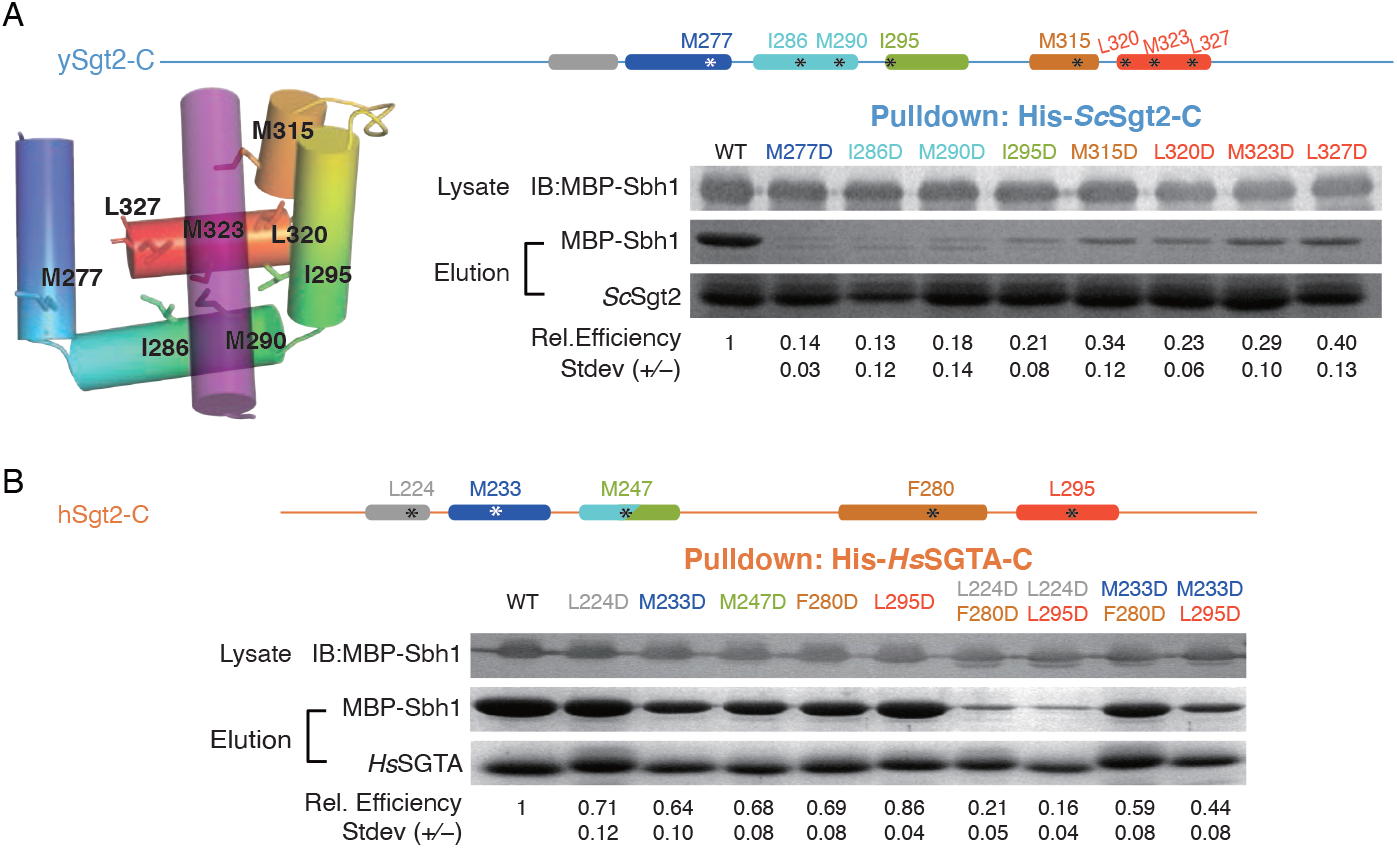
Effects on TA client binding of charge mutations to the putative hydrophobic groove of Sgt2-C_cons_. For these experiments, individual point mutations are introduced into Sgt2-C and tested for their ability to capture Sbh1 quantified as in Figure 2*D*. (*A*) For ySgt2-C, a schematic and cartoon model are provided highlighting the helices and sites of individual point mutants both color-ramped for direct comparison. For the cartoon, the docked TMD is shown in purple. Binding of MBP-Sbh1 to his-tagged ySgt2-C and mutants were examined as in Figure 2*D*. Lanes for mutated residues are labeled in the same color as the schematic *(B)* Same analysis as in *A* for hSgt2-C. In addition, double point mutants are included.

### Sgt2-C domain binds clients with a hydrophobic segment ≥ 11 residues

With a molecular model for ySgt2-C_cons_ and multiple lines of evidence for a hydrophobic groove, we sought to better understand the specific requirements for TMD binding. We and others have demonstrated that a monomeric C-domain from Sgt2 is sufficient for binding to TA clients [13]. To study the minimal constraints on TA client binding, we chose to focus on a monomeric construct of Sgt2 (Sgt2-TPR-C) binding to variable TMDs. TA clients were designed where the overall (sum) and average (mean) hydrophobicity, length, and the distribution of hydrophobic character were varied in the TMDs. These artificial TMDs, a Leu/Ala helical stretch followed by a Trp, were constructed as C-terminal fusions to the soluble protein BRIL (Fig. 7*A*). The total and mean hydrophobicity are controlled by varying the helix-length and the Leu/Ala ratio. For clarity, we define a syntax for the various artificial TA clients to highlight the various properties under consideration: hydrophobicity, length, and distribution. The generic notation is TMD-length[number of leucines] which is represented, for example, as 18[L6] for a TMD of 18 amino acids containing six leucines.

**Fig. 7.**
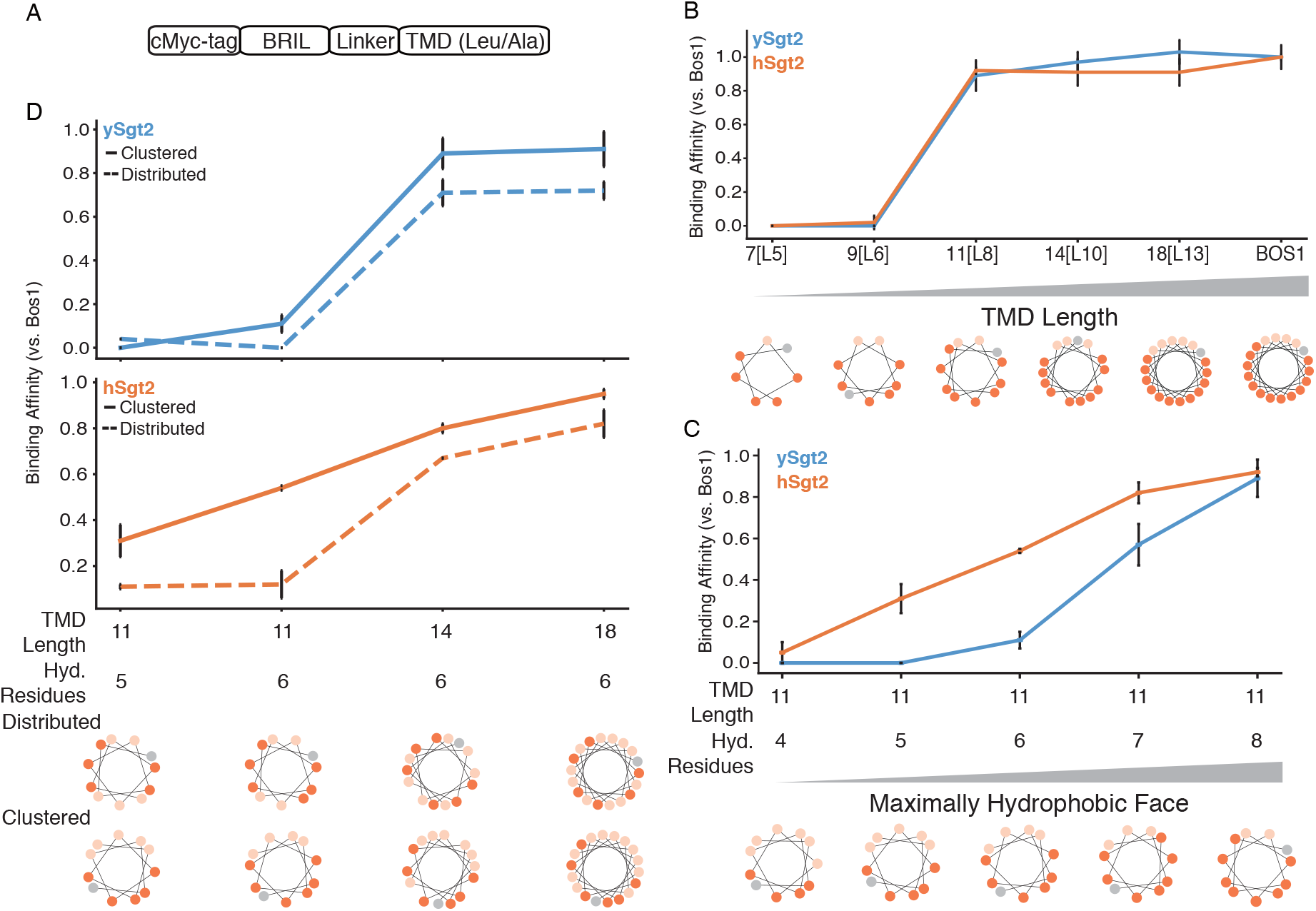
Minimal requirements for client recognition by Sgt2. (*A*) Schematic of model TA clients. Quantification of complex formation is calculated and normalized to that of complexes with Bos1-TMD, here defined as relative binding efficiency. (*B*) Complex formation of ySgt2 (blue) and hSgt2 (orange) with the TA client Bos1_TMD_ and several artificial TA clients noted x[Ly], where x denotes the length of the TMD and y denotes the number of leucines in the TMD. The helical wheel diagrams of the TMD of TA clients here and for subsequent panels with leucines colored in dark orange, alanines colored in pale orange, and tryptophans colored in grey. (*C*) Complex formation of ySgt2-TPR-C and hSgt2-TPR-C with artificial TA clients with TMDs of length 11 and increasing numbers of leucine. (*D*) Comparison of complex formation of ySgt2-TPR-C and hSgt2-TPR-C with artificial TA clients of the same lengths and hydrophobicities but differences in the distribution of leucines, i.e. clustered (solid line) vs distributed (dotted line).

Our first goal with the artificial clients was to define the minimal length of a TMD to bind to the C-domain. As described earlier, captures of his-tagged Sgt2-TPR-C with the various TA clients were performed. We define a relative binding efficiency as the ratio of captured TA client by a Sgt2-TPR-C normalized to the ratio of a captured wild-type TA client by Sgt2-TPR-C. In this case we replaced the TMD in our artificial clients with the native TMD of Bos1 (Bos1_TMD_). The artificial client 18[L13] shows a comparable binding efficiency to Sgt2-TPR-C as that of Bos1_TMD_ (Fig. 7*B*). From the helical wheel diagram of the TMD for Bos1, we noted that the hydrophobic residues favored one face of the helix. We explored this ‘hydrophobic face’ by using model clients that maintained this orientation while shortening the length and maintaining the average hydrophobicity of 18[L13] (Fig. 7*B*). Shorter helices of 14 or 11 residues, 14[L10] and 11[L8], also bound with similar affinity to Bos1. Helices shorter than 11 residues, 9[L6] and 7[L5], were not able to bind Sgt2-TPR-C (Fig. 7*B*) establishing a minimal length of 11 residues for the helix consistent with the dimensions of the groove predicted from the structural model (Fig. 3). In the context of the full-length Sgt2, which exists as a dimer, an 11-residue cut-off suggests that two C-domains could come together and bind to a single TA whose TMD lengths range from 18-24 residues.

Since a detected binding event occurs with TMDs of at least 11 amino acids, we decided to probe this limitation further. The dependency of client hydrophobicity was tested by measuring complex formation of Sgt2-TPR-C and artificial TA clients containing an 11 amino acid TMD with increasing number of leucines (11[Lx]). As shown in Fig. 7*C*, increasing the number of leucines monotonically enhances complex formation, echoing previous results [56]. hSgt2-TPR-C binds to a wider spectrum of hydrophobic clients than ySgt2-TPR-C, which could mean it has a more permissive hydrophobic binding groove, also reflected in the milder impact of Ala replacement and Asp mutations in hSgt2-TPR-C to TA client binding (Fig. 2*C* and Fig. 6*A*).

### Sgt2-C preferentially binds to TMDs with a hydrophobic face

Next, we address the properties within the TMD of TA clients responsible for Sgt2 binding. In the case of ySgt2, it has been suggested that the co-chaperone binds to TMDs based on hydrophobicity and helical propensity [56]. In our system, our artificial TMDs consist of only alanines and leucines which have high helical propensities [57], and despite keeping the helical propensity constant and in a range that favors Sgt2 binding, there is still variation in binding efficiency. For the most part, varying the hydrophobicity of an artificial TA client acts as expected, the more hydrophobic TMDs bind more efficiently to Sgt2 TPR-C (Fig. 7*C*). Our C_cons_ model suggests the hydrophobic groove of ySgt2-C protects a TMD with highly hydrophobic residues clustered to one side (see Fig. 3*B*). To test this, various TMD pairs with the same hydrophobicity, but different distributions of hydrophobic residues demonstrates TA clients with clustered leucines have a higher relative binding efficiency than those with a more uniform distribution (Fig. 7*D*). Helical wheel diagrams demonstrate the distribution of hydrophobic residues along the helix (*e.g.* bottom Fig. 7*D*). The clustered leucines in the TMDs create a hydrophobic face which potentially interacts with the hydrophobic groove formed by the Sgt2-C_cons_ region, corresponding to the model in Fig. 3*B*.

## Discussion

Sgt2, the most upstream component of the GET pathway, plays a critical role in the targeting of TA IMPs to their correct membranes. Its importance as the first confirmed selection step of ER versus mitochondrial [56] destined TA IMPs necessitates a molecular model for TA client binding. Previous work demonstrated a role for the C-domain of Sgt2 to bind to hydrophobic clients, yet the exact binding domain remained to be determined. Through the combined use of biochemistry, bioinformatics, and computational modeling, we conclusively identify the minimal client-binding domain of Sgt2. Pulldown experiments revealed a minimal six α-helical region as the TMD binding domain of Sgt2 that is also conserved in structure and sequence analyses. This domain preferentially binds TMDs with hydrophobic residues organized onto one face of the helix. While NMR spectroscopy displays an intrinsically disordered domain in the absence of substrate, computational models of ySgt2-C consistently predict a structured helical hand in its conserved region, while the rest of the domain remained flexible and varied between models. Structural similarities between the ySgt2-C model and the STI1 domains DP1 and DP2 now place Sgt2 among a class of co-chaperones. Together, these results allow us to present a validated structural model of the Sgt2 C-domain as a methionine-rich helical hand for grasping a hydrophobic helix and to provide a mechanistic explanation for binding a TMD of at least 11 hydrophobic residues.

We confidently identify the C-domain of Sgt2 as containing a STI1 domain for client binding through sequence alignments and structural homology. This places Sgt2 into a larger context of conserved co-chaperones (Fig. 8*A*). In the co-chaperone family, the STI1 domains predominantly follow HSP-binding TPR domains connected by a flexible linker, reminiscent of the domain architecture of Sgt2. As noted above, for Sti1 these domains are critical for client-processing and coordinated hand-off between Hsp70 and Hsp90 homologs [58] as well as coordinating the simultaneous binding of two heat shock proteins. Both Sgt2 and the co-chaperone Hip coordinate pairs of TPR and STI1 domains by forming stable dimers via their N-terminal dimerization domains [59]. With evidence for a direct role of the carboxylate-clamp in the TPR domain of Sgt2 for client-binding now clear [21], one can speculate that the two TPR domains may facilitate TA client entry into various pathways that use multiple heat shock proteins.

**Fig. 8.**
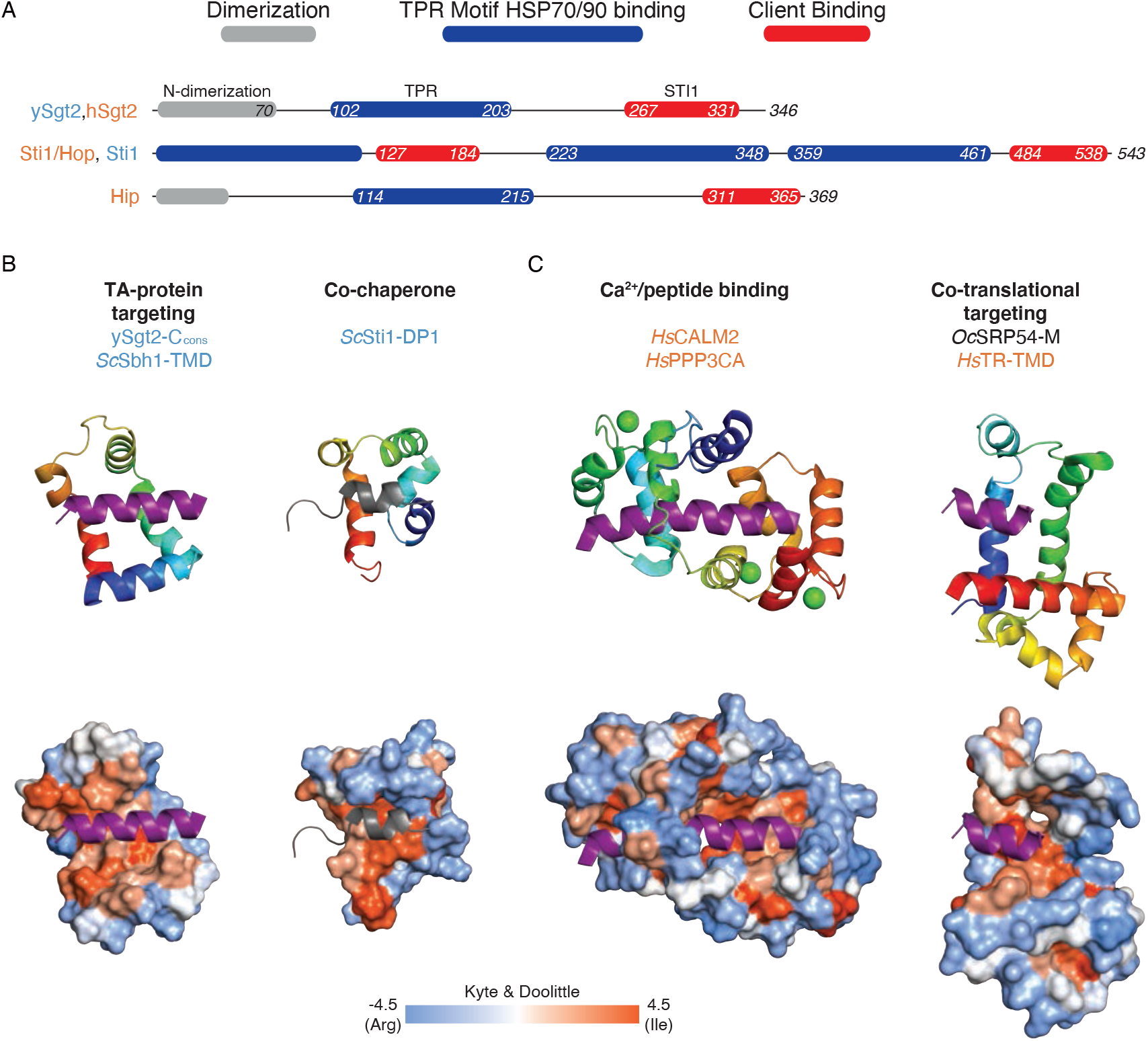
Various domain structures of STI1 and other helical-hand containing proteins. (*A*) The domain architectures of proteins with a STI1 domain were obtained initially from InterPro [87] and then adjusted as discussed in the text. Each domain within a protein is colored relative to the key. (*B*) Structural comparison of various hydrophobic-binding helical-hand protein complexes. For each figure only relevant domains are included. Upper row, color-ramped cartoon representation with bound helices in purple. Lower row, accessible surface of each protein colored by hydrophobicity again with docked helical clients in purple. In order, the predicted complex of ySgt2-C_cons_ and *Sc*Sbh2-TMD, DP1 domain from yeast Sti1 with N-terminus containing H0 in grey (*Sc*Sti1-DP1)(PDBID: 2LLV), human calmodulin (*Hs*CALM2) bound to a hydrophobic domain of calcineurin (*Hs*PPP3CA) (PDBID: 2JZI), and M domain of SRP54 from *Oryctolagus cuniculus* (*Oc*SRP54-M) and the signal sequence of human transferrin receptor (*Hs*TR-TMD) (PDBID: 3JAJ).

Computational modeling reveals a conserved region sufficient for TA client binding that consists of a helical hand of five alpha-helices that form a hydrophobic groove to bind the client TMD. The concept of TMD binding by a helical hand is reminiscent of other proteins involved in membrane protein targeting. Like Sgt2, the signal recognition particle (SRP) contains a methionine-rich domain that binds signal sequences and TMDs. While the helical order is inverted, again five amphipathic helices form a hydrophobic groove that cradles the client signal peptide [60] (Fig. 8*B*). Here once more, the domain has been observed to be flexible in the absence of client [61,62] and, in the resting state, occupied by a region that includes a helix which must be displaced [60]. Another helical-hand example recently shown to be involved in TA IMP targeting is calmodulin where a crystal structure reveals two helical hands coordinating to clasp a TMD at either end (Fig. 8*B*) [63]. Considering an average TMD of 18-20 amino acids (to span a ~40Å bilayer), each half of calmodulin interacts with about 10 amino acids. This is in close correspondence to the demonstrated minimal 11 amino acids for a TA client to bind to the monomeric Sgt2-TPR-C. In the context of the full-length Sgt2, one can speculate that the Sgt2 dimer may utilize both C-domains to bind to a full TMD, similar to calmodulin. Cooperation of the two Sgt2 C-domains in client-binding could elicit conformational changes in the complex that could be recognized by downstream factors, such as additional interactions that increase the affinity to Get5/Ubl4A.

Intriguingly, Sgt2-TPR-C preferentially binds to artificial clients with clustered leucines. If the C-domain forms a hydrophobic groove as suggested by the computational model, it provides an attractive explanation for this preference. In order to bind to the hydrophobic groove, a client buries a portion of its TMD in the groove leaving the other face exposed. Clustering hydrophobic residues contributes to the hydrophobic effect driving binding efficiency and protecting them from the aqueous environment. Indeed, GET pathway substrates have been suggested to be more hydrophobic TMDs than EMC substrates [64]. Of these, for the most hydrophobic substrates, like Bos1, residues on both sides of the TMD could be protected by a pair of C-domains. Alternatively, the unstructured N-terminal loop through H0 could act as a lid surrounding the circumference of the client’s TMD. Unstructured regions participating in substrate binding as well as capping a hydrophobic groove have both suggested in the context of other domains, *e.g.* with Get3 [4]. The role for this clustering of hydrophobic residues in TA client recognition and targeting merits further investigation.

What is the benefit of the flexible helical-hand structure for hydrophobic helix binding? While it remains an open question, it is notable that evolution has settled on similar simple solutions to the complex problem of specific but temporary binding of hydrophobic helices. For all of the domains mentioned above, the flexible helical-hands provide an extensive hydrophobic surface to capture a client-helix—driven by the hydrophobic effect. Typically, such extensive interfaces are between pairs of pre-ordered surfaces resulting in high affinities and slow off rates. These helical hands are required to only engage temporarily, therefore the flexibility offsets the favorable free energy of binding by charging an additional entropic cost for ordering a flexible structure in the client-bound complex. The benefit for TA clients is a favorable transfer to downstream components in the GET pathways as seen for ySgt2 [21] and hSgt2 [50]. The demonstration that TA transfer from hSgt2 to Get3 is twice as fast as disassociation from hSgt2 into solution, perhaps interaction with Get3 leads to conformational changes that further favor release [50].

While hSgt2 and ySgt2 share many properties, there are a number of differences between the two homologs that may explain the different biochemical behavior. For the C_cons_-domains, hSgt2 appears to be more ordered in the absence of client as the peaks in its NMR spectra are broader (Fig. 1*E*). Comparing the domains at the sequence level, while the high glutamine content in the C-domain is conserved it is higher in hSgt2 (8.8% versus 15.2%). The additional glutamines are concentrated in the predicted longer H4 helix (Fig. 1*A*). The linker to the TPR domain is shorter compared to ySgt2 while the loop between H3 and H4 is longer. Do these differences reflect different roles? As noted, in every case the threshold for hydrophobicity of client-binding is lower for hSgt2 than ySgt2 (Fig. 1*E*, 5, and 6) implying that the mammalian protein is more permissive in client binding. The two C-domains have similar hydrophobicity, so this difference in binding might be due to a lower entropic cost paid by having the hSgt2 C-domain more ordered in the absence of client or the lack of an unstructured N-terminal loop.

The targeting of TA clients presents an intriguing and enigmatic problem for understanding the biogenesis of IMPs. How subtle differences in each client modulates the interplay of hand-offs that direct these proteins to the correct membrane remains to be understood. In this study, we focus on a central player, Sgt2 and its client-binding domain. Through biochemistry and computational analysis, we provide a structural model that adds more clarity to client discrimination.

## Material and Methods

### Plasmid constructs

MBP-Sbh1 full length, *ySgt2*_95-346_ (*ySgt2*-TPR-C), *ySgt2*_222-346_ (*ySgt2*-C), *ySgt2*_260-327_ (*ySgt2*-C_cons_), *ySgt2*_266-327_ (*ySgt2*-ΔH0), *hSgt2*_87-313_ (*hSgt2*-TPR-C), *hSgt2*_213-313_ (*hSgt2*-C), *hSgt2*_219-300_ (*hSgt2*-C_cons_), and *hSgt2*_228-300_ (*hSgt2*-ΔH0) were prepared as previously described [12,65]. Genes of *ySgt2* or *hSgt2* variants were amplified from constructed plasmids and then ligated into an pET33b-derived vector with a 17 residue N-terminal hexa-histidine tag and a tobacco etch virus (TEV) protease site. Single or multiple mutations on Sgt2 were constructed by site-direct mutagenesis. Artificial TA clients were constructed in a pACYC-Duet plasmid with a N-terminal cMyc tag, BRIL fusion protein [66], GSS linker, and a hydrophobic C-terminal tail consisting of leucines and alanines and ending with a tryptophan.

### Protein expression and purification

All proteins were expressed in *Escherichia coli* NiCo21 (DE3) cells (New England BioLabs). To co-express multiple proteins, constructed plasmids were co-transformed as described [65]. Protein expression was induced by 0.3 mM IPTG at OD_600_ ~ 0.7 and harvested after 3 hours at 37°C. For structural analysis, cells were lysed through an M-110L Microfludizer Processor (Microfluidics) in lysis buffer (50 mM Tris, 300 mM NaCl, 25 mM imidazole supplemented with benzamidine, phenylmethylsulfonyl fluoride (PMSF), and 10 mM Δ-mercaptoethanol (BME), pH 7.5). For capture assays, cells were lysed by freeze-thawing 3 times with 0.1 mg/mL lysozyme. To generate endogenous proteolytic products of *ySgt2*-TPR-C for MS analysis, PMSF and benzamidine were excluded from the lysis buffer. His-tagged Sgt2 and his-tagged Sgt2/TA complexes were separated from the lysate by batch incubation with Ni-NTA resin at 4°C for 1hr. The resin was washed with 20 mM Tris, 150 mM NaCl, 25 mM imidazole, 10 mM BME, pH 7.5. The complexes of interest were eluted in 20 mM Tris, 150 mM NaCl, 300 mM imidazole, 10 mM BME, pH 7.5.

For structural analysis, the affinity tag was removed from complexes collected after the nickel elution by an overnight TEV digestion against lysis buffer followed by size-exclusion chromatography using a HiLoad 16/60 Superdex 75 prep grade column (GE Healthcare).

Measurement of Sgt2 protein concentration was carried out using the bicinchoninic acid (BCA) assay with bovine serum albumin (BSA) as standard (Pierce Chemical Co.). Samples for NMR and CD analyses were concentrated to 10-15 mg/mL for storage at −80°C before experiments.

For the *in vitro* transfer assay, plasmids encoding for the full-length ySgt2 cysteine mutants were transformed into BL21 Star cells (Invitrogen). Cells were grown in 2x yeast-tryptone (2xYT) media and induced with 0.1mM IPTG at an OD_600_ of 0.6 then harvested after 3 hours at 30°C by centrifugation. Cells were lysed in 50mM Tris pH 8.0, 500mM NaCl, 10% glycerol, and 1x BugBuster (Millipore Sigma), supplemented with protease inhibitors (4-(2-aminoethyl)benzenesulfonyl fluoride hydrochloride (Roche), benzamidine, and BME). Full-length his-tagged ySgt2 and cysteine mutants were separated from the lysate by batch incubation with Ni-NTA resin (Qiagen) at 4°C for 1 hour. The resin was washed with 50mM Tris pH 8.0, 500mM NaCl, 10% glycerol, and 25mM imidazole and then the protein was eluted in 50mM Tris pH 8.0, 500mM NaCl, 10% glycerol, and 300mM imidazole. For storage, protein was dialyzed in 25mM K-HEPES pH 7.5, 150mM KOAc, and 20% glycerol at 4°C and then flash frozen in liquid nitrogen. Purified Bos1 with p-benzoyl-l-phenylalanine (BPA) labeled at residue 230 (Bos1_BPA_) and yeast Ssa1 were gifts from the lab of Shu-ou Shan (Caltech).

### NMR Spectroscopy

_15_N-labeled proteins were generated from cells grown in auto-induction minimal media as described [67] and purified in 20 mM phosphate buffer, pH 6.0 (for ySgt2-C, 10mM Tris, 100mM NaCl, pH 7.5). The NMR measurements of _15_N-labeled Sgt2-C proteins (~0.3-0.5 mM) were collected using a Varian INOVA 600 MHz spectrometer at either 25°C (ySgt2-C) or 35°C (hSgt2-C) with a triple resonance probe and processed with TopSpin™ 3.2 (Bruker Co.).

### CD Spectroscopy

The CD spectra were recorded at 24°C with an Aviv 202 spectropolarimeter using a 1 mm path length cuvette with 10 μM protein in 20 mM phosphate buffer, pH 7.0. The CD spectrum of each sample was recorded as the average over three scans from 190/195 to 250 nm in 1 nm steps. Each spectrum was then decomposed into its most probable secondary structure elements using BeStSel [68].

### Glu-C digestion of the double cysteine mutants on *ySgt2-C*

Complexes of the co-expressed wild type or double cysteine mutated His-*ySgt2*-TPR-C and the artificial TA client, 11[L8], with either a cMyc or MBP tag were purified as the other His-Sgt2 complexes described above or initially purified via amylose affinity chromatography before nickel chromatography explained earlier. The protein complexes were mixed with 0.2 mM CuSO_4_ and 0.4 mM 1,10-phenanthroline at 24°C for 20 min followed by 50 mM N-ethyl maleimide for 15 min. Sequencing-grade Glu-C protease (Sigma) was mixed with the protein samples at an approximate ratio of 1:30 and the digestion was conducted at 37°C for 22 hours. Digested samples were mixed with either non-reducing or reducing SDS-sample buffer, resolved via SDS-PAGE using Mini-Protean® Tris-Tricine Precast Gels (10-20%, Bio-Rad), and visualized using Coomassie Blue staining.

### *In vitro* transfer assay of Bos1 from Ssa1 to *ySgt2*

The *in vitro* transfer assays were performed as in Chio et al. 2019 and Shao et al. 2017 [49,50]. Specifically, 39μM Bos1_BPA_ (50mM HEPES, 300mM NaCl, 0.05% LDAO, 20% glycerol) was diluted to a final concentration of 0.1μM and added to 4μM Ssa1 supplemented with 2mM ATP (25mM HEPES pH7.5, 150mM KOAc). After one minute, 0.3μM of full-length ySgt2 or mutant was added to the reaction. Samples were flash frozen after one minute and placed under a 365nm UV lamp for 2 hours on dry ice to allow for BPA crosslinking.

### Protein immunoblotting and detection

For western blots, protein samples were resolved via SDS-PAGE and then transferred onto nitrocellulose membranes by the Trans-Blot® Turbo™ Transfer System (Bio-Rad). Membranes were blocked in 5% non-fat dry milk and hybridized with antibodies in TBST buffer (50 mM Tris-HCl pH 7.4, 150 mM NaCl, 0.1% Tween 20) for 1 hour of each step at 24°C. The primary antibodies were used at the following dilutions: 1:1000 anti-penta-His mouse monoclonal (Qiagen), 1:5000 anti-cMyc mouse monoclonal (Sigma), and a 1:3000 anti-Strep II rabbit polyclonal (Abcam). A secondary antibody conjugated either to alkaline phosphatase (Rockland, 1:8000) or a 800nm fluorophore was employed, and the blotting signals were chemically visualized with either the nitro-blue tetrazolium/5-bromo-4-chloro-3′-indolyphosphate (NBT/BCIP) chromogenic assay (Sigma) or infrared scanner. All blots were photographed and quantified by image densitometry using ImageJ [69] or ImageStudioLite (LI-COR Biosciences).

### Quantification of Sgt2-TA complex formation

The densitometric analysis of MBP-Sbh1 capture by His-Sgt2-TPR-C quantified the intensity of the corresponding protein bands on a Coomassie Blue G-250 stained gel. The quantified signal ratios of MBP-Sbh1/His-Sgt2-TPR-C are normalized to the ratio obtained from the wild-type (WT). Expression level of MBP-Sbh1 was confirmed by immunoblotting the MBP signal in cell lysate. Average ratios and standard deviations were obtained from 3-4 independent experiments.

In artificial TA client experiments, both his-tagged Sgt2-TPR-C and cMyc-tagged artificial TA clients were quantified via immunoblotting signals. The complex efficiency of Sgt2-TPR-C with various TA clients was obtained by

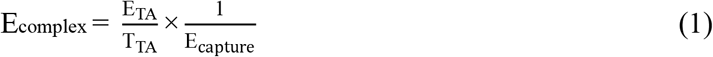

where E_TA_ is the signal intensity of an eluted TA client representing the amount of TA client co-purified with Sgt2-TPR-C and T_TA_ is the signal intensity of a TA client in total lysate which corresponds to the expression yield of that TA client. Identical volumes of elution and total lysate from different TA clients experiments were analyzed and quantified. In order to correct for possible variation the amount of Sgt2-TPR-C available for complex formation, E_capture_ represents the relative amount of Sgt2-TPR-C present in the elution (E_Sgt2_) compared to a pure Sgt2-TPR-C standard (E_purified,Sgt2_).

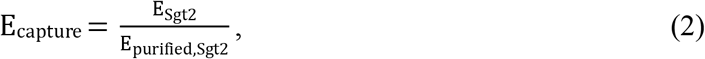

Each E_TA_ and T_TA_ value was obtained by blotting both simultaneously, *i.e.* adjacently on the same blotting paper. To facilitate comparison between TA clients, the Sgt2-TPR-C/TA client complex efficiency E_complex,TA_ is normalized by Sgt2-TPR-C/Bos1 complex efficiency E_complex,Bos1_.

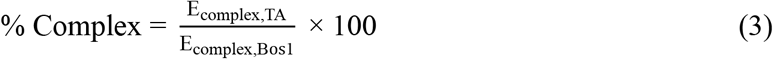

### Sequence alignments

An alignment of Sgt2-C domains was carried out as follows: all sequences with an annotated N-terminal Sgt2/A dimerization domain (PF16546 [70]), at least one TPR hit (PF00515.27, PF13176.5, PF07719.16, PF13176.5, PF13181.5), and at least 50 residues following the TPR domain were considered family members. Putative C-domains were inferred as all residues following the TPR domain, filtered at 90% sequence identity using CD-HIT [71], and then aligned using MAFFT G-INS-i [72]. Other attempts with a smaller set (therefore more divergent) of sequences results in an ambiguity in the relative register of H0, H1, H2, and H3 when comparing Sgt2 with SGTA.

Alignments of Sti1 (DP1/DP2) and STI1 domains were created by pulling all unique domain structures with annotated STI1 domains from Uniprot. Where present, the human homolog was selected and then aligned with PROMALS3D [73]. PROMALS3D provides a way of integrating a variety of costs into the alignment procedure, including 3D structure, secondary structure predictions, and known homologous positions.

All alignments were visualized using Jalview [74].

### Molecular modeling

Putative models for ySgt2-C were generated with I-TASSER, PCONS, Quark, Robetta (*ab initio* and transform-restrained modes), Phyre2, and RaptorX via their respective web servers [48,75–78]. The highest scoring model from Quark was then chosen to identify putative TA client binding sites by rigid-body docking of various transmembrane domains modelled as α-helices (3D-HM [79]) into the ySgt2-C_cons_ through the Zdock web server [80]. Pairwise distances were calculated between C_β_ atoms (the closer C_α_ proton on glycine) using mdtraj [81]. Based on our disulfide crosslinks, new models were predicted using Robetta in *ab initio* mode specifying C_β_-C_β_ atom distance constraints bounded between 0 and 9 Å.

For hSgt2, using the same set of structure prediction servers above, we were only able to produce a clear structural model using the Robetta transform-restrained mode. We were also unable to generate a reliable model by directly using the ySgt2-C model as a template [82]. To crosslink distance data from ySgt2 as restraints for hSgt2, pair positions were transferred from one protein to the other via an alignment of Sgt2-C domains (excerpt in Fig. 1A) and ran Robetta *ab initio*. Also, we grafted the N-terminal loop of ySgt2-C on hSgt2-C with the same set of restraints.

Images were rendered using PyMOL 2.3 (www.pymol.org).

## Acknowledgements

We thank D. G. VanderVelde for assistance with NMR data collection; S. Mayo for providing computing resources; S. Shan, H. J. Cho, Y. Liu, and members of the Clemons lab for support and discussion. We thank J. Mock and A. M. Thinn for comments on the manuscript. This work was supported by the National Institutes of Health (NIH) grants GM105385 and GM097572 (to WMC), NIH/National Research Service Award Training Grant GM07616 (to SMS and MYF), and a National Science Foundation Graduate Research fellowship Grant 1144469 (to SMS).

---

**Fig. S1.**
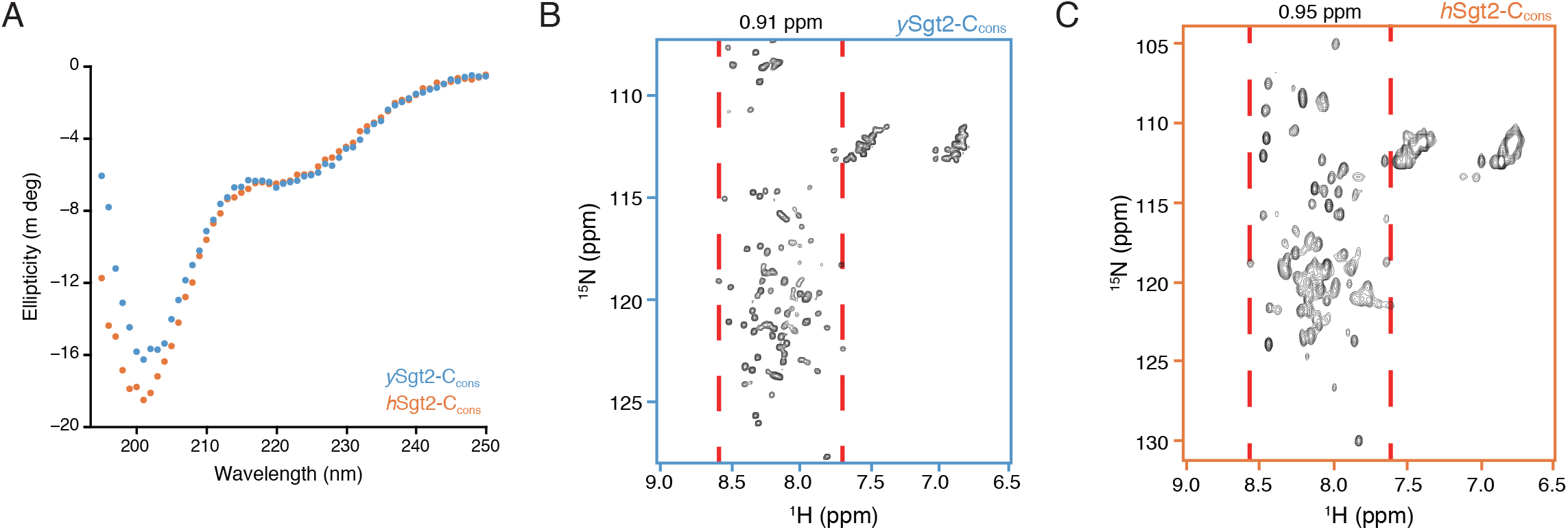
Biophysical characterization of the Sgt2-C_cons_ domain. (*A*) CD spectra as in Fig. 1*C* for the conserved C-terminal domains of ySgt2 (blue) and hSgt2 (orange). NMR spectra as in Fig. 1*D* & *E* for ySgt2-C_cons_ (*B*, blue) and hSgt2-C_cons_ (*C*, orange).

**Fig. S2.**
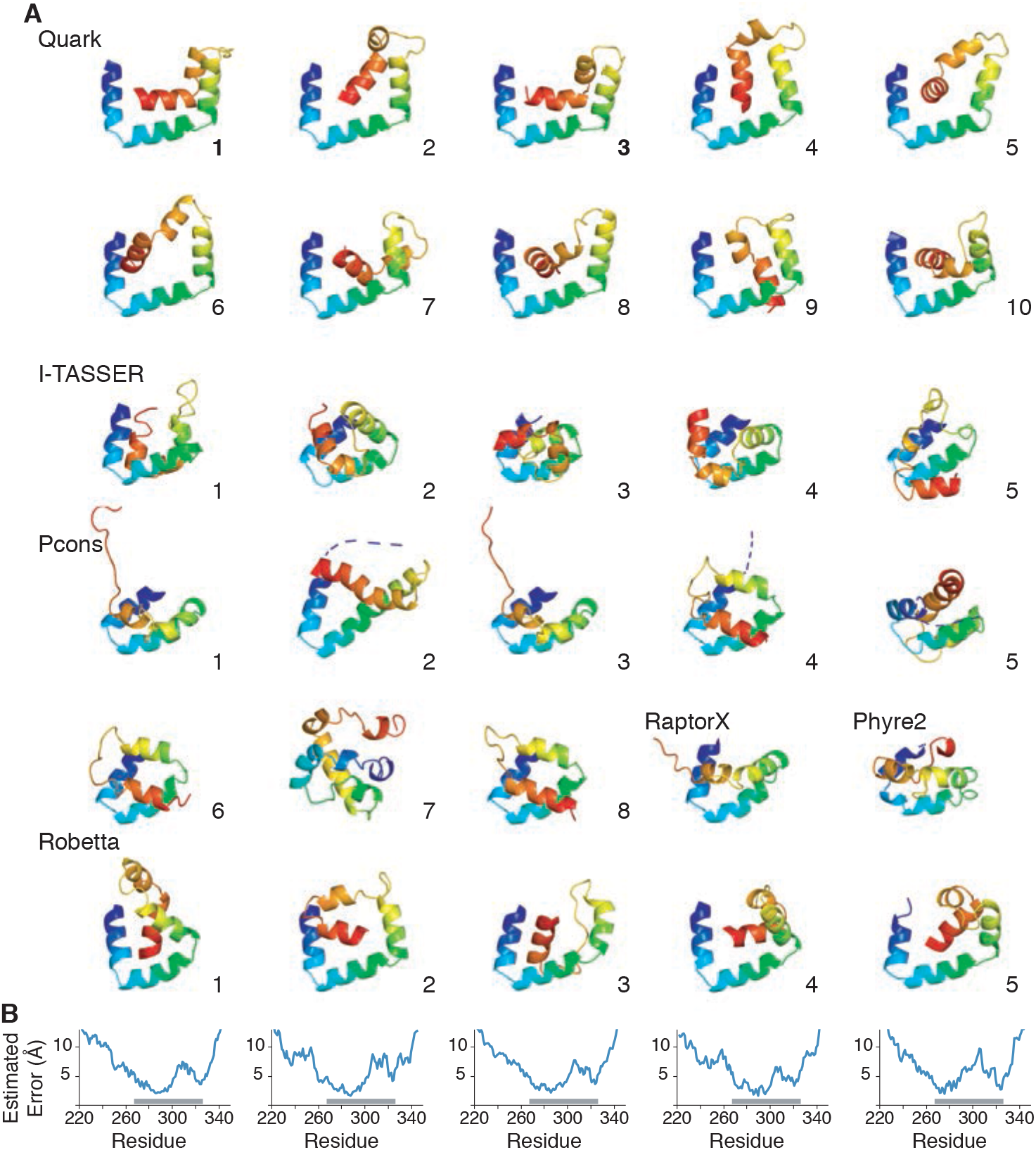
Structural models across prediction methods. (*A*) Predictions from Quark, I-TASSER, Pcons, Phyre2, RaptorX, and Robetta. Methods produce between 5 and 10 models. (*B*) Robetta provides a residue-wise estimated error in Angstroms; this is shown below the corresponding models with a grey bar indicating the C_cons_ region.

**Fig. S3.**
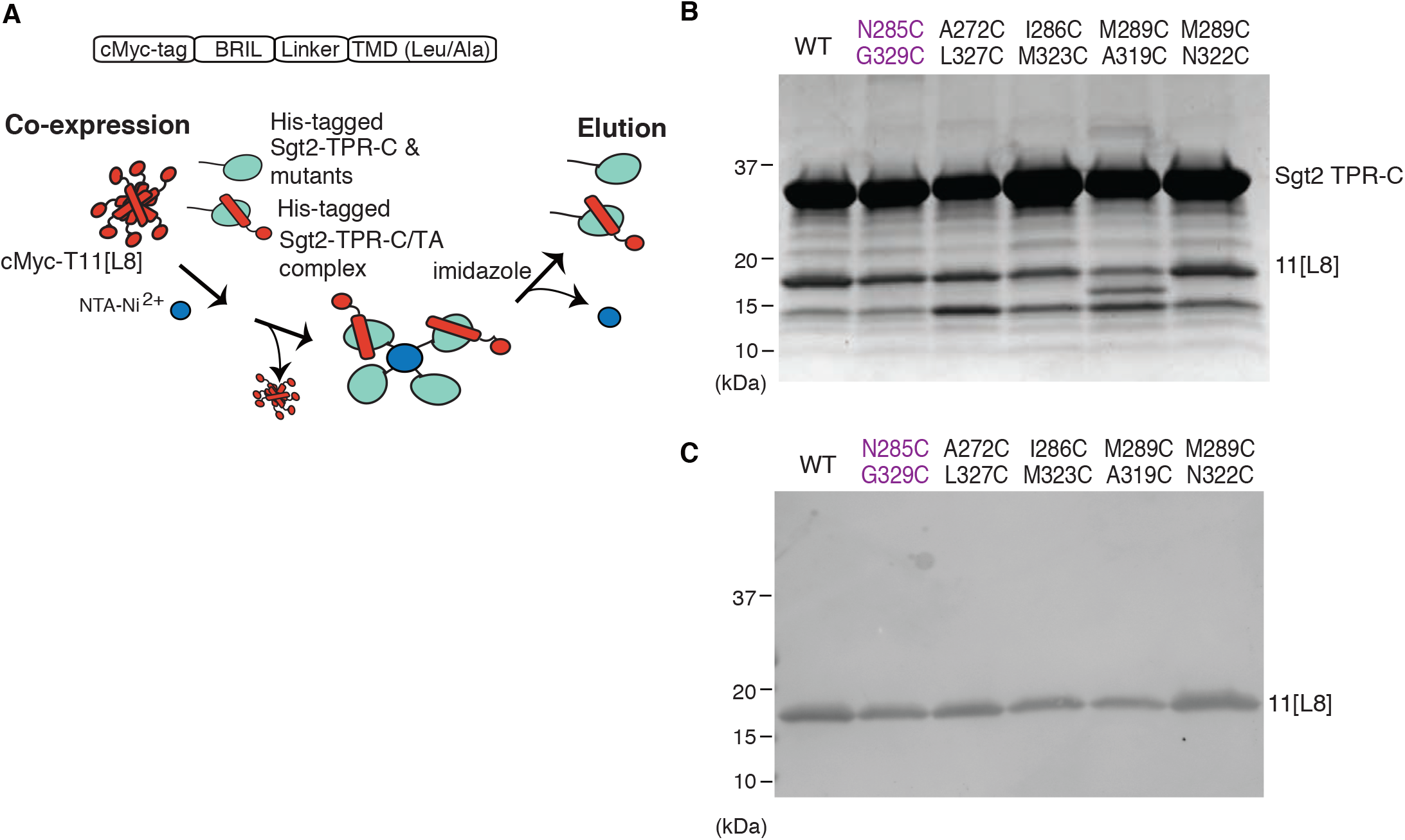
Cysteine mutants are capable of binding to TA clients. (*A*) Schematic showing how his-tagged ySgt2-TPR-C and double cysteine mutant constructs were coexpressed with the TA client 11[L8], and complexes were purified by nickel affinity chromatography. (*B*) A SDS-PAGE gel of the elution fractions demonstrates that 11[L8] was present in the elution suggesting double cysteine mutations do not affect client binding. (*C*) An anti-cMyc western blot of the fractions represented in the SDS-PAGE gel also demonstrates that 11[L8] was present in all elutions.

**Fig. S4.**
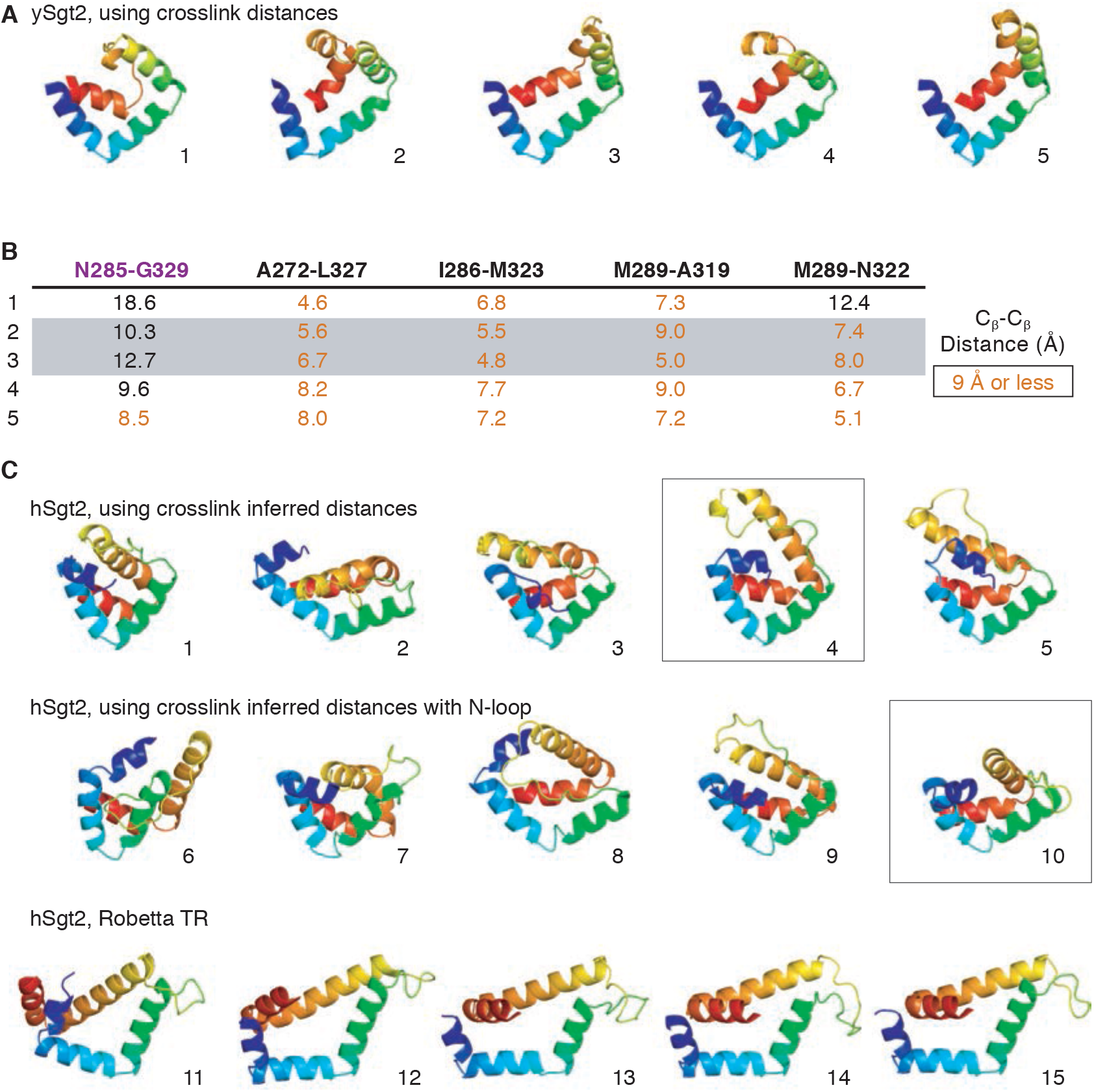
Distance restraints lead to improved ySgt2-C and suggestive hSgt2-C models. (*A*) Prediction of ySgt2-C using distances from in vitro crosslinking. (*B*) C_β_-C_β_ distances between residues probed by *in vitro* disulfide crosslinking for each ySgt2 model. Distances 9 Å or less are colored orange. For models where all distances correspond (4 near and 1 far), the row is shaded grey. (*C*) Models for hSgt2-C using restraints, adding a N-terminal loop, and via the new Robetta TR method.

